# Humans can navigate complex graph structures acquired during latent learning

**DOI:** 10.1101/723072

**Authors:** Milena Rmus, Harrison Ritz, Lindsay E Hunter, Aaron M Bornstein, Amitai Shenhav

**Affiliations:** Department of Psychology, University of California, Berkeley; Department of Cognitive, Linguistic, and Psychological Sciences, Brown University; Department of Psychology, Princeton University; Department of Cognitive Sciences, University of California, Irvine; Center for the Neurobiology of Learning and Memory, University of California, Irvine; Carney Institute for Brain Science, Brown University

## Abstract

Humans appear to represent many forms of knowledge in associative networks whose nodes are multiply connected, including sensory, spatial, and semantic. Recent work has shown that explicitly augmenting artificial agents with such *graph-structured* representations endows them with more human-like capabilities of compositionality and transfer learning. An open question is how humans acquire these representations. Previously, it has been shown that humans can learn to navigate graph-structured conceptual spaces on the basis of direct experience with trajectories that intentionally draw the network contours (Schapiro et al., 2012;2016), or through direct experience with rewards that covary with the underlying associative distance (Wu et al., 2018). Here, we provide initial evidence that this capability is more general, extending to learning to reason about shortest-path distances across a graph structure acquired across disjoint experiences with randomized edges of the graph - a form of latent learning. In other words, we show that humans can *infe*r graph structures, assembling them from disordered experiences. We further show that the degree to which individuals learn to reason correctly and with reference to the structure of the graph corresponds to their propensity, in a separate task, to use *model-based* reinforcement learning to achieve rewards. This connection suggests that the correct acquisition of graph-structured relationships is a central ability underlying forward planning and reasoning, and may be a core computation across the many domains in which graph-based reasoning is advantageous.

## Introduction

Humans have a remarkable ability to construct representations of their environment by integrating sparse observations, and to use these representations to control different forms of complex behavior (Whittington et al., 2020). These sorts of structure representations allow us to plan the words we will use to communicate a new idea; plan a route through an unfamiliar city; or plan an event months or even years away. Acquiring such representations allows for efficient learning, and facilitates the transfer of learned strategies between environments with similar structures (Sutton, Precup & Singh,1999). So far, research has mainly focused on how individuals learn by linking states that are experienced in temporal sequences (Schapiro et al., 2012; Schapiro et al., 2016). However, experiences in the real world are rarely ordered in this way. Rather, people often have to infer these underlying structures representations from sparse or disconnected experiences. How they achieve this remains unclear. Here, we combine a graph theoretic approach with a novel task to examine how people infer the structure of their environments under such conditions of sparsity, and then test for links between one’s ability to infer structure under these conditions, and their use of structure information when engaging in goal-directed planning.

### Inferring structure

One body of work has examined how people develop internal models of their environment based on their experience with individual states in that environment and the transitions between them (Fermin et al. 2010; Behrens et al. 2018). Foundational research in the area demonstrated that animals construct *cognitive maps* as they navigate their spatial environment (Tolman, 1948; O’Keefe and Nadel, 1978), and that neural representations of these maps (decoded from regions of hippocampus) not only reflect the animal’s location in that space but also (1) their recent locations and (2) the future projections of locations they intend to visit (Johnson & Redish, 2007). Recent work has shown that cognitive maps can also be extrapolated from *abstract* learned associations. For instance, Schapiro and colleagues (2013) built a virtual graph-like structure, with each node represented by an individual abstract stimulus. In their experiment, participants traversed this graph sequentially, one node at a time. Despite never seeing the underlying graph, participants were able to recover the graph, based on their experience of the likelihood of moving from one node to another. In addition, much like the representation of spatial maps, the graph representation itself could also be decoded from their brain activity. Similar forms of construction and navigation have been demonstrated over episodic and semantic memory representations, including connections between words (Jurafsky, 1996), concepts (Collins & Quillian 1969), events (Collin et al. 2015), and people (Parkinson et al., 2017; Tamir & Thornton, 2018; FeldmanHall & Shenhav, 2019).

However, for the most part, research in this area has studied how people develop cognitive models of their environment in contexts where they are transitioned sequentially through that environment, eliminating the need for the individual to infer relationships between states that are not directly linked. These studies therefore fail to capture a distinctive property of human cognition -- the ability to infer latent structure based on sparse (e.g. non-sequential) information (Bunsey and Eichenbaum, 1996). For instance, simulations by Whittington and colleagues (2020) recently demonstrated that agents do not need to experience all edges connecting nodes in a graph network to demonstrate the knowledge of the graph structure. Rather, just visiting different parts of the graph (nodes) allows for near-optimal transitive inference performance. The model developed by these researchers (Tolman-Eichenbaum machine) captures these properties, and proposes a detailed account of the neural mechanisms that enable such robust learning features (combination of grid cells and place cell remapping). Hence, it is crucial to also probe cognitive inference processes in a task that omits temporal structure, and requires individuals to integrate information from disjoint experiences.

### Goal-directed planning

The advantages of building internal models by inference extend beyond just navigation benefits. A separate body of work has examined the process by which people navigate these internal models in order to determine the course of action that will maximize their future rewards (Sutton and Barto 1998; Daw et al. 2005; Daw et al. 2011). Early work demonstrated that these goal-directed forms of decision-making trade off against habitual behavior, enabling an animal to adapt to rapid changes in their environment (Balleine & Dickinson 1998; Balleine & O’Doherty, 2009). More recently, it has been shown that goal-directed decision-making can be formalized as *model-based reinforcement learning* (RL) (Daw et al., 2005; Sutton and Barto,1998; Dolan & Dayan, 2013). Model-based RL represents a form of RL that stores a model of how different states in the environment are connected to one another (e.g., the likelihood of transitioning from one state to another), and the rewards that an agent can expect upon reaching a given state. By contrast, *model-free* forms of RL only store the value of previous actions taken in a given state, and therefore are less sensitive to changes in the structure of one’s environment (e.g., if a certain state is no longer rewarded or if two states are no longer connected, requiring a detour).

These two types of RL - model-based and model-free - are commonly dissociated with a task developed by Daw and colleagues (2011). In this task, participants must choose between a pair of states at 2 sequential stages, at the end of which they receive an outcome (either rewarding or non-rewarding). Importantly, the 2 stages are linked via probabilistic transitions, and the probability of receiving the reward at the second stage states gradually shifts during the task. Therefore, in order to behave in a goal-directed way, though not necessarily earn more points, one must integrate information about transitions and the outcomes. The patterns of decisions on this “two-step task” can therefore reveal the extent to which a participant engages in *model-free* decision-making - choosing actions based only on whether they were recently rewarded - or *model-based* planning - choosing actions based on a consideration of both the recent rewards and the likelihood of reaching those rewards given the transition structure of the task environment (Daw et al., 2011; Decker et al., 2016). Using this task, researchers have shown that individual differences in one’s tendency to engage in model-based decision-making have been linked to variability in working memory capacity (Otto et al. 2013), cognitive control (Daw, Niv & Dayan 2005; Otto et al. 2015), temporal discounting (Shenhav, Rand & Greene 2017; Hunter, Bornstein, Hartley 2019) and psychiatric symptoms associated with compulsive behavior and social isolation (Gillan et al., 2016).

### Inferential ability as a potential constraint on goal-directed planning

Goal-directed planning thus depends critically on both our ability to (1) *learn the structure of one’s environment and* (2) our ability to *leverage the representation of this structure in pursuit of rewards*. Recently, a consensus has developed that these capacities share overlapping computational and neural substrates (Collin et al. 2015; Shohamy & Turk-Browne 2013; Behrens et al. 2018; Vikbladh et al. 2019). However, while the mechanisms that support learning, navigating, and deploying an internal model have separately been well-characterized, the relationships between these domains remain poorly understood. In particular, work on model-based planning uses tasks in which the associative structure is made explicit (Daw et al., 2011; Konovalov & Krajbich; 2016; 2020),de-emphasizing structure learning and leaves behavioral measures of goal-directed behavior to index how these representations are used. As a result, little is known about whether and how one’s ability to infer the structure of an environment relates to their ability to leverage such a representation when engaging in goal-directed planning. Recent work hints that when provided with more information about the task structure (i.e. more detailed instructions), participants appear more goal-directed/model-based (da Silva and Hare, 2020). This suggests that having information about the structure boosts behavioral patterns consistent with model-based planning, providing more empirical evidence of the link between model-based planning and structure representations.

In the current work, we developed a novel set of tasks to measure participants’ ability to infer the structure of an abstract (non-spatial) graph, based on disjoint experiences with pairs of adjacent nodes throughout that graph. Multiple measures supported the hypothesis that participants are able to infer the structure of the graph by integrating over sparse information. Choices and response times revealed that participants were able to flexibly reason about relative and overall distances within the graph, and an explicit reconstruction task demonstrated that their representation of graph structure preserved metric distance information. We also asked whether the measures indexing structure inference correlated with measures of model-based planning from a separate task. We found that participants who exhibited better structure inference ability were also more likely to engage in model-based planning in the two-step task. This work validates a novel approach to measuring individual differences in the ability to infer and navigate latent structures in one’s environment, while also providing preliminary evidence of connection between structure inference and model-based planning.

## Methods

### Participants

We recruited 81 participants (38 female, Age range 18-27, Mean(SD) Age = 20(1.8)) from the Brown University participant pool. Participants received either course credit or monetary compensation for participating in the study. All participants provided informed consent in accordance with the policies of the Brown University Institutional Review Board. We excluded two participants with a high rate of perseverative responses in the two-step task (repeating the same response on more than 95% of the trials), and 2 participants due to the issues with data saving, resulting in the sample of total 77 participants included in the analyses. This study was approved by the Brown University Institutional Review Board.

### Procedure

#### Two-Step Task

The two-step task is a sequential decision-making task, which enables assessment of dissociation between model-based and model-free choice strategies. In the task, participants made choices on two sequential stages, with the aim of obtaining a reward. In this version of the task (Decker et al, 2016), participants chose between two spaceships at stage one, which probabilistically transitioned to one of the two states (planets) at stage two (*Figure 1A*). In particular, each of the spaceships transitioned to one of the planets 70% of the time (common transition), and to the other planet 30% of the time (rare transition). These transition probabilities remained fixed throughout the task, and were taught to participants during training. At the second stage, participants encountered two aliens and chose one to solicit the space treasure/reward. The two aliens awarded treasure with independent probability, which shifted slowly over time according to a Gaussian random walk. Participants were instructed to earn as many pieces of treasure as possible. They had 3 seconds to make their choice on each stage. If they failed to make a response within the given time frame, a red ‘x’ appeared on top of the stimulus, and the trial terminated. Participants performed 40 practice trials, followed by 200 experimental trials.

**Figure 1.**
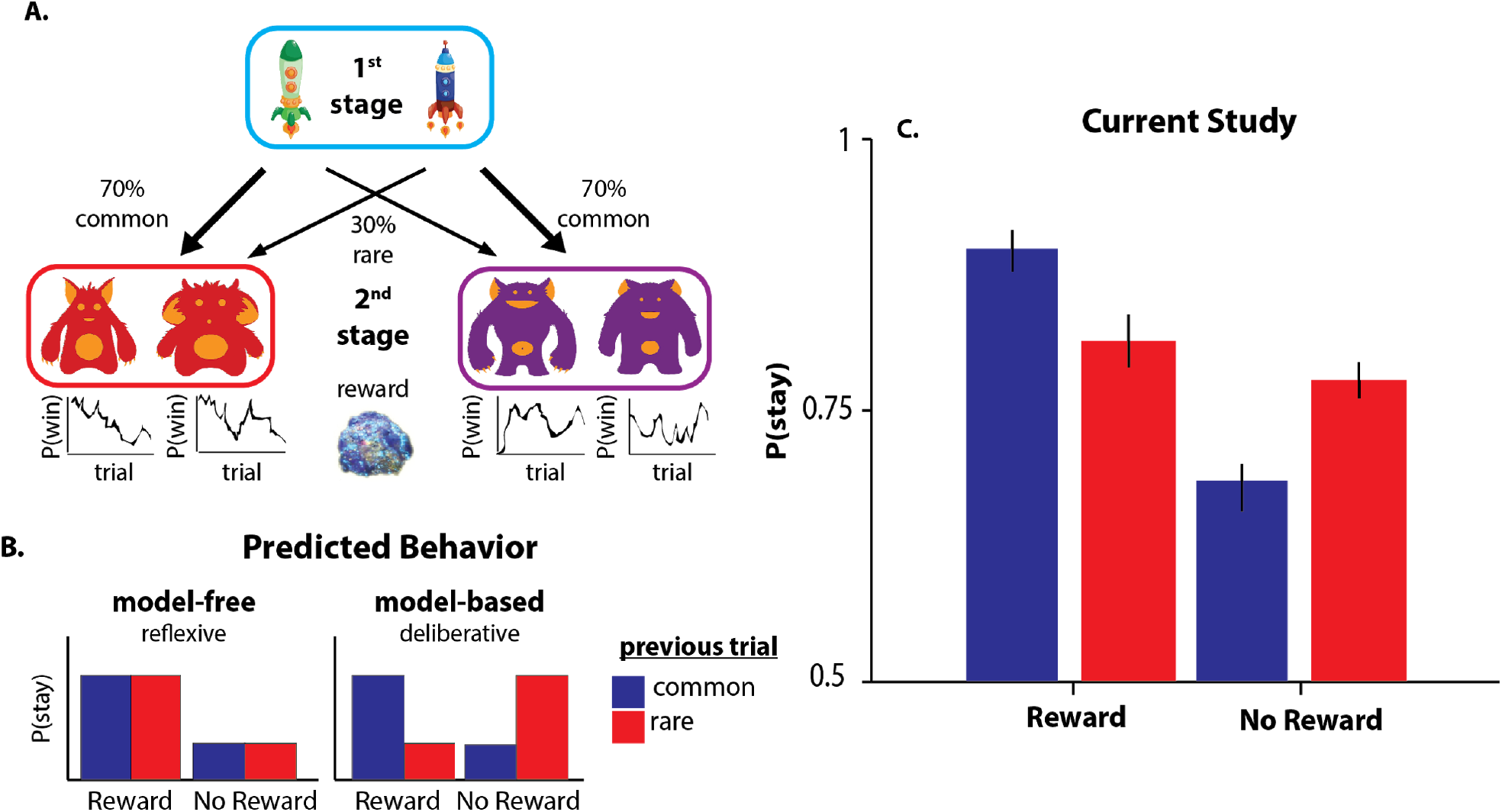
**A)** Two-step task design, adapted from Decker et al. (2016). The fixed structure of the probabilistic transitions from 1^st^ stage states 2^nd^ stage states enables the distinction of model-based and model-free choices by examining the influence of the previous trial on the subsequent first-stage choice. **B)** A model-free learner tends to repeat previously rewarded first-stage choices (“stay”), regardless of the transition type that led to the reward (a main effect of reward on subsequent first-stage choices). By contrast, a model-based learner exploits knowledge of the transition structure and will favor the first-stage action that is most likely to lead to the same state if rewarded and the action least likely to lead to the same state if not rewarded (a reward-by-transition interaction effect on subsequent first-stage choices). **C)** Consistent with previous literature, our results show that the participants exhibit a mixture of model-based and model-free choice strategies.

The two-step task characterizes dissociable trial-by-trial adjustment of stage 1 choices, reflecting model-free and model-based choice strategies. On each trial, participants could choose between repeating the previous stage 1 choice and switching to the other spaceship. The model-free strategy predicts that the likelihood of staying or switching (repeating or changing the previous choice) on the first stage is informed by the outcome of the previous trial. The model-based strategy, on the other hand, predicts that the arbitration between staying and switching based on the observed outcome is modulated by the knowledge of transition type (common or rare) which on average led to that outcome over the course of trials. Thus, model-free reasoners choose to stay (repeat their prior stage 1 choice) if the outcome of that choice was rewarding on the previous trial, regardless of the transition type. On the other hand, model-based reasoners utilize the transition structure to select options that will most likely transition to the rewarding state (*Figure 1B*).

#### Graph task

Following the two-step task, participants performed a structure inference task designed to assess their ability to infer the latent structure based on the sequence of disjoint node pairs which, when reassembled, form the graph. They viewed a sequence of object pairs, each of which represented a pair of adjacent nodes drawn at random from an underlying undirected graph with 12 nodes and 16 edges (*Figure 2A*). Nodes in the graph were tagged by images of objects, which were randomly assigned for each participant. Each node-pair was presented for 1 second on the screen, after which the trial terminated and the next pair was presented.

**Figure 2.**
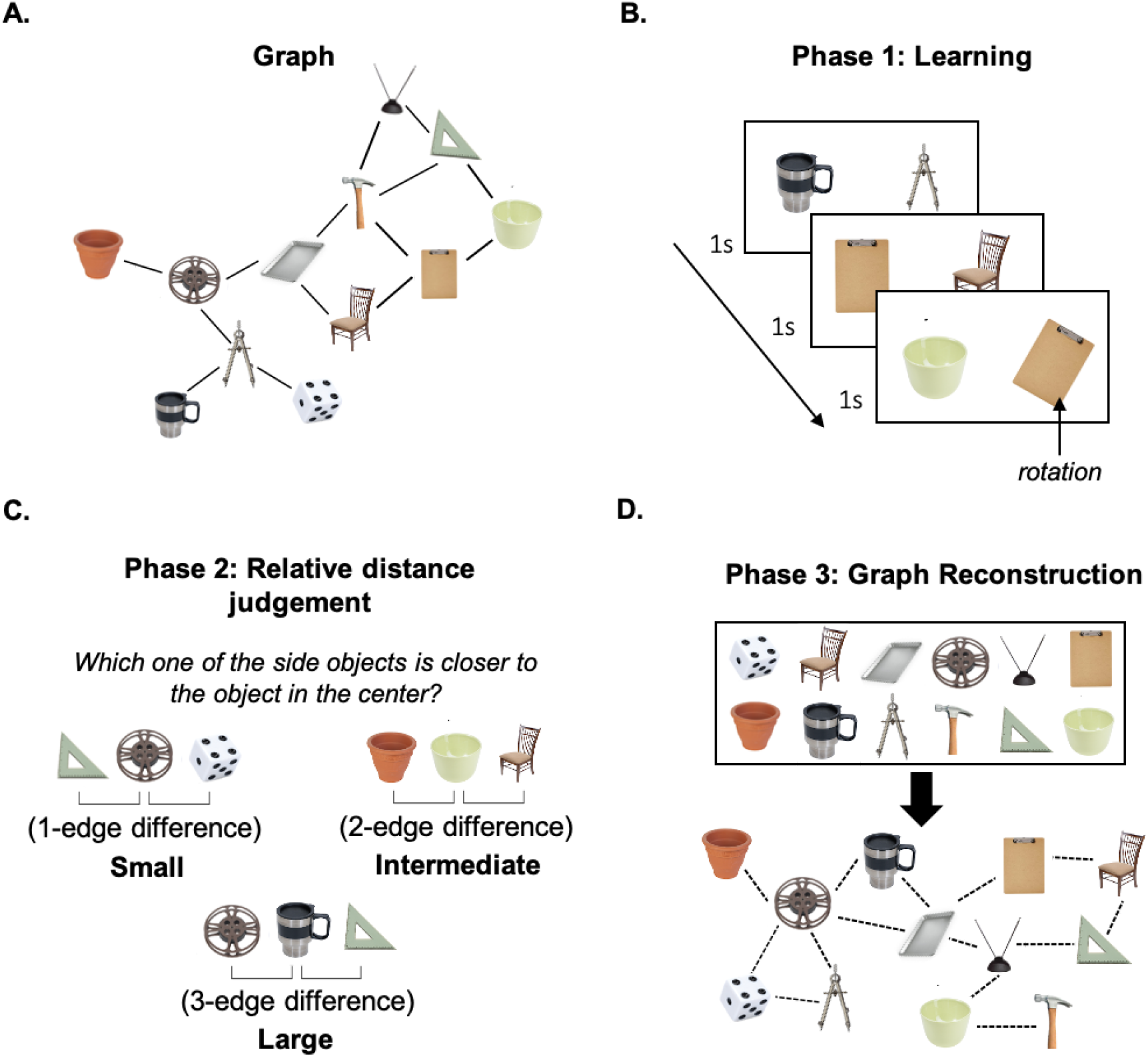
Schematic of the graph task. **A)** Underlying graph. Node labels (the object images) were fully randomized across all subjects. **B)** Learning phase: Participants observed pairs of randomly sampled adjacent nodes. **C)** Relative distance judgment task trial: participants were asked to identify the more proximal node. **D)** Graph reconstruction: participants freely arranged objects and connections between them, based on the learned object associations.

#### Phase 1: Learning

In phase 1, participants passively viewed a sequence of object pairs, each displaying a pair of adjacent nodes. Node pairs were drawn at random, such that consecutive trials sampled adjacent node pairs from different locations on the graph (*Figure 2B*). Participants were not informed that there was an underlying structure, but were told that they would be tested on their memory of the pairs that had been presented. Each of the pairs was presented 44 times. In order to ensure that participants sustained attention throughout this phase, they were also asked to respond any time an object in the pair was rotated from its default position (which occurred on 10% of trials).

#### Phase 2: Relative distance judgment task

In phase 2, participants performed a relative distance judgment task, which required them to judge the relative distance between three randomly-selected nodes, unconstrained by edge relationship. In particular, participants viewed two nodes on either side of the screen and were asked to indicate which was closer to the reference node (shown centrally) based on the pairs they had seen in Phase 1 (*Figure 2C*). Participants were presented with 204 trials, and had an unlimited amount of time to make their choice.

#### Phase 3: Graph reconstruction

In phase 3, participants were shown all 12 nodes they had encountered in Phases 1 and 2 (*Figure 2D*). They were instructed to arrange the objects freely, by using their mouse to click on and move the object images on the screen. Once they positioned the nodes on the screen, participants were asked to connect them by clicking on pairs of images that they wished to group together. Participants had an unlimited amount of time to complete this stage of testing, and were allowed to make as many connections as they wished.

All of the tasks were programmed in Matlab (version 2016b; Natick, Massachusetts: The MathWorks Inc), using the Psychtoolbox 3 extension (Brainard, 1997; Pelli, 1997; Kleiner et al, 2007).

### Analysis

#### Two-step task

Following previous work (Gillan, Otto, Phelps, & Daw, 2015; Daw et al, 2011; Decker et al, 2015), we quantified model-based behavior by performing a logistic mixed-effects regression analysis. We modeled participants’ choice to stay (repeat the previous stage 1 choice) or switch on the current trial, as a function of (1) previous outcome (reward or no reward) (2) transition type (common or rare), and (3) a reward-by-transition type interaction (*Stay ~ Previous Reward * Transition Type + ( 1+Previous Reward * Transition Type | Participant*)). The main effect of the reward in the model is an index of model-free behavior, quantifying participant’s choices as a function of recent outcome. The reward-by-transition interaction term serves as an index of model-based behavior, as it captures how much participants’ choices were affected by the recent outcome, modulated by their knowledge of the transition structure. Therefore, variability in the interaction term demonstrates individual differences in how much participants relied on model-based reasoning in the task. The regression included maximal random slopes and intercepts for each participant.

#### Reinforcement Learning Model

Participants’ full trial-by-trial choice sequence in the task can also be fit with a computational reinforcement-learning model that gauges the degree to which participants’ choices are better described by a model-based or model-free reinforcement learning algorithm. Indices of model-based learning in the two-step RL task were derived via Bayesian estimation using a variant of the computational model introduced in (Daw et al., 2011). The model assumes choice behavior arises as a combination of model-free and model-based reinforcement learning. Each trial *t* begins with a first-stage choice *c*_1,*t*_ followed by a transition to a second state *s_t_* where the participant makes a 2^nd^ stage choice *c*_2,*t*_ and receives reward *r_t_*. Upon receipt of reward *r_t_*, the expected value of the chosen 2^nd^ stage action (the left vs. the right alien) 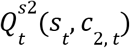 is updated in light of the reward received.

According to the model, the decision-maker uses a learned value function over states and choices *Q*^*s*2^(*s, c*) to makes second-stage choices. On each trial, the value estimate for the chosen action is adjusted towards the reward received using a simple delta rule, 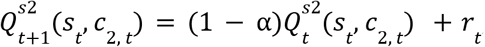, where α is a free learning rate parameter that dictates the extent to which value estimates are updated towards the received outcome on each trial. Unlike the standard delta rule 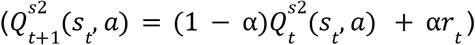, in this equation and in similar references throughout, the learning rate α is omitted from the latter term. Effectively, this reformulation rescales the magnitudes of the rewards by a factor of 1/α and the corresponding weighting (e.g., temperature) parameters β by α. The probability of choosing a particular 2^nd^ stage action *c*_2,*t*_ in state *s_t_* is approximated by a logistic softmax, 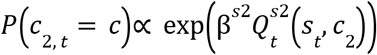, with free inverse temperature parameter β^*s*2^ normalized over both options *c*_2_.

First-stage choices are modeled as a product of both model-free and model-based value predictions. The model-based value of each 1^st^ stage choice is dictated by the learned value of the corresponding 2^nd^ stage state, maximized over the two actions: 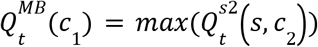, where *s* is the second-stage state predominantly produced by first-stage choice *c*_1_. Model-free values are governed by two learning rules, TD(0) and TD(1), each of which updates according to a delta rule towards a different target. Whereas earlier models posit a single model-free choice weight β_*MF*_ and use an eligibility trace parameter λ∈(0, 1) to control the relative contributions of TD(0) and TD(1) learning, here, as in recent work by Gillan et al., (2016), model-free valuation is split into its component TD(0) and TD(1) stages, each with separate sets of weights and Q values. TD(0) backs-up the value of the stage-1 choice on the most recent trial 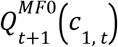 with the value of the state-action pair that immediately (e.g., lag-0) followed it: 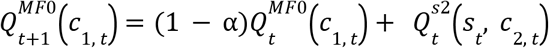. TD(1), on the other hand, backs up its value estimate 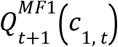 by looking an additional step ahead (e.g., lag-1) at the reward received at the end of the trial: 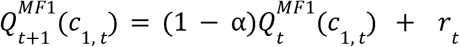. Ultimately, Stage-1 choice probabilities are given by a logistic softmax, where the contribution of each value estimate is weighted by its own model free temperature parameter:

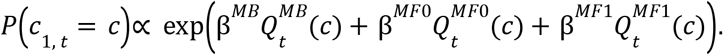

At the end of each trial, the value estimates for all unchosen actions and unvisited states are multiplicatively re-weighted by a free discount parameter γ [0,1]. This conventional parameterization reflects the assumption that value estimates decay exponentially at a rate of 1 − γ over successive trials (Ito & Doya, 2009; Hunter et al., 2018). The temporal decay of value is widely endorsed by normative and empirical research on reinforcement learning (Sutton & Barto, 1998; Ito & Doya, 2009), and is further motivated endogenously by the perseverative nature of choice behavior in this task. Earlier models of behavior in this task have operationalized choice perseveration using a “stickiness” parameter, which is implemented as a recency bonus or subjective “bump” in the value of whichever first stage action was chosen on the most recent trial (irrespective of reward) (Daw et al., 2011). Including a decay parameter also accounts for the fact that people tend to ‘stay’ (repeat) their previous 1^st^ stage choice: as γ approaches 0, the value of the unchosen actions decreases relative to the value of the chosen action on the next trial regardless of whether or not a reward was received (Hunter et al., 2018). In total the model has six free parameters: four weights (β^*s*2^, β^*MB*^, β^*MF*0^, β^*MF*1^), a learning rate α, and a decay rate γ. The six free parameters of the model (β^*s*2^, β^*MB*^, β^*MF*0^, β^*MF*1^, α, γ) were estimated by maximizing the likelihood of each individual’s sequence of choices. Numerical optimization was used to find maximum likelihood estimates of the free parameters, with 10 random initializations to help avoid local optima.

#### Graph task

We first assessed whether choice difficulty predicts accuracy and RTs in the relative distance judgment task. We defined choice difficulty as the absolute difference between exemplars’ distances from the reference node, with distance defined as the shortest path length. The greater the distance difference between options and the reference, the closer one option was to the reference relative to the other, thus making the discrimination easier. In addition, we tested whether participants’ response times also scaled with the total distance of option nodes from the reference node (e.g. depth-first search). As with the regression approach in the two-step task, we computed random effects for all subjects in these regressions as well.

We also quantified how similar each participant’s recovered graph was to the true graph using two metrics. First, we looked at the average similarity between the adjacency matrix of the ground truth and the recovered graph (*Figure 5A*). Second, we tested whether we can predict the pairwise ground truth distance (the shortest path) with the Euclidean distance between any two node placements (*Figure 5B*.)

#### Relationship between the performance on the graph task and the two-step task

To answer the question of whether there is a relationship between variability in structure inference and the planning task, we looked at the relationship between the model-based indices and six main measures from the graph task: 1) overall judgment accuracy, random effect estimates of task difficulty on 2) response times and 3) accuracy, 4) effect of total distance on the search time, 5) percentage of correctly identified edges, 6) Fisher-transformed pixel-node distance correlations. We added these separate metrics from the graph task as covariates in the 1-back *stay ~ reward * transition* logistic regression model, in order to estimate direct association between model-based planning and these between-subject factors. We report results from independent models, with each z-scored covariate *Z* entered separately: *stay ~ reward * transition type * Z + (1+reward*transition type|Participant)*. In addition, using principal component analysis (PCA) we collapsed the six graph measures into a single latent factor which captures variability in structure inference performance. We then performed the same analysis described above, where we entered participants’ PCA scores as z-scored covariates in the two-step task regression model. To validate consistency between our two approaches to extracting subject-level model-based planning indices from the two step task (the interaction term from the logistic regression, and the model-based weighting parameter (*β*_MB_) from the computational model), we also tested the association between participants’ PCA scores and *β*_MB_ by performing a robust linear regression and Spearman correlation.

## Results

Participants (N=77) performed a novel structure inference and judgment task with three main phases (*Figure 2*). In Phase 1, participants were given the opportunity to implicitly learn a graph-like structure through experiences with pairs of nodes in that graph. On each of 704 trials, they viewed a pair of objects (e.g., a bowl and a clipboard) and asked to report whether one of the objects was rotated relative to a canonical orientation (*Figure 2B*). Though they were never informed of this, each of these object pairs reflected a randomly drawn pair of adjacent nodes from an underlying graph, for which each node was represented by a single object (e.g., a clipboard) (*Figure 2A*). In Phase 2, participants made a series of judgments about the relative distances of three randomly selected nodes in the graph. On each trial, they were asked to evaluate which of two objects they thought was “closer” to a third reference object (*Figure 2C*), requiring them to make implicit inferences about the latent structure of the graph. In Phase 3, participants were asked to freely arrange the 12 objects and their connections in order to reconstruct their best estimate of the underlying graph (*Figure 2D*).

### Evidence of structure inference

In Phase 1, participants performed very well on the rotation detection task (mean accuracy = 89%, SD = 5%; d’ = 2.65, SD = .8; false alarm rate= 0.5%), indicating that they sustained attention throughout the learning phase. We then examined two sets of measures to assess whether participants were able to infer the underlying graph structure based on this learning phase.

First, using our relative distance judgment task (Phase 2), we tested how fast and how accurate they were at judging how far two objects are from one another within the underlying graph. Despite only having experienced disjoint node pairs from this graph and never having been made explicitly aware of the graph itself, participants were significantly above chance in discerning distances between nodes they had never seen paired together (mean accuracy = 67%, SD = 13%; *t(76)* = 11.49, *p* = 2.5e-18). Importantly, these distance judgments were also sensitive to the overall difficulty of the distance judgment: participants were both faster (*Figure 3*; *β* =−.03 (.01), *t(76)* =−2.30, *p* = .02) and more accurate (*Figure 3*; *β* =.02 (.008), *t(76)* = 3.50, *p* = .0004) the closer the reference was to the target, relative to the foil. Response times thus appeared tp reflect the relative time it might have taken participants to search for one node relative to the other (from the reference node). If that were the case, RTs should not only scale with the relative distance between the nodes, but also with the *total* distance from the reference to the two other nodes (reflecting the depth of search). Consistent with this prediction, participants’ response times were longer when the total distance of both options to the target was greater (*β* =.13, *t(76)* =7.07, *p* = 5.9e-10).

**Figure 3.**
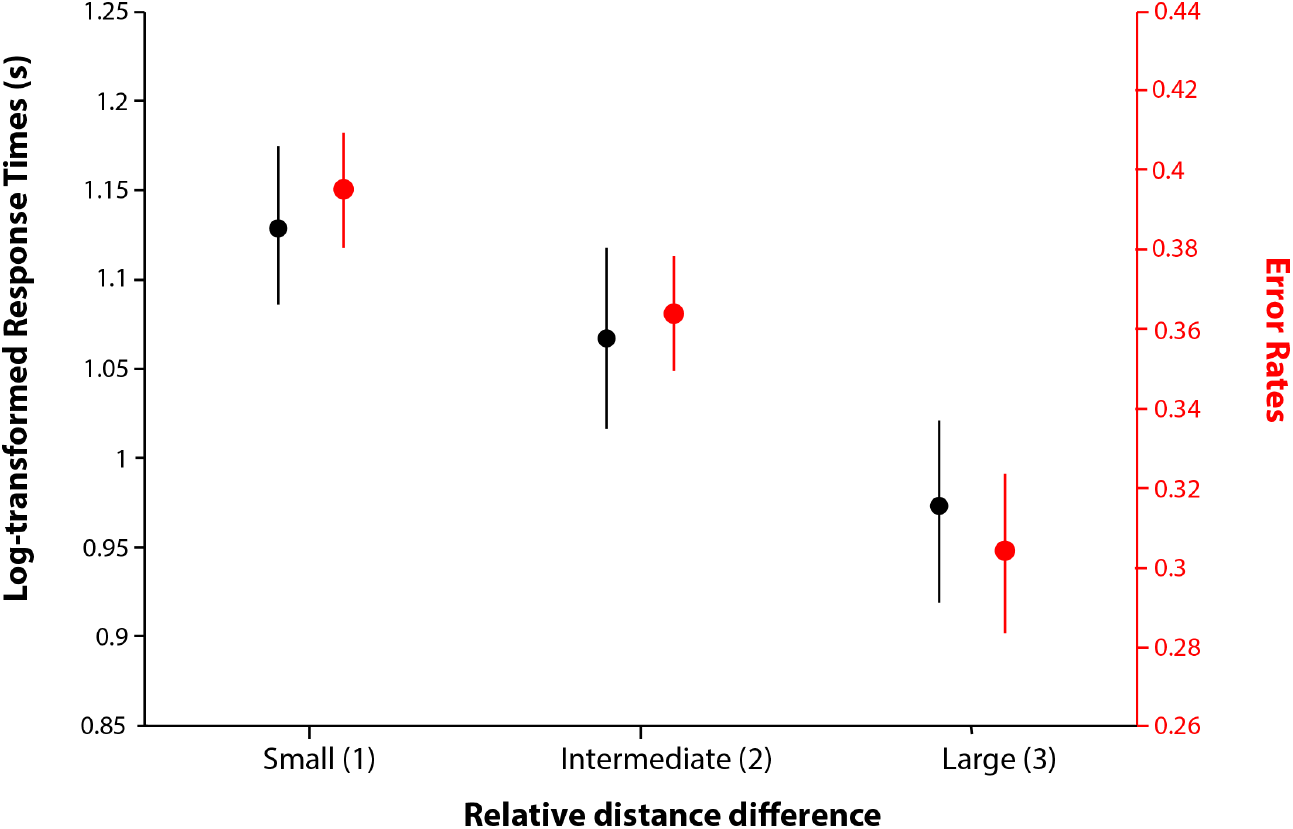
Participants are faster (black) and more accurate (red) at selecting the closer node when it was much closer than the alternative. Relative distance is discretized for display purposes.

These patterns of accuracy and RT as a function of node distance suggest that participants were able to implicitly infer the structure of the underlying graph. We extended these analyses to further understand *how* participants represent this structure (Figure 4A). For instance, it is possible that participants used a global representation of the distances between nodes (e.g., a map-like representation; Behrens et al. 2018), which should allow for an efficient search that only depends on the route between nodes and targets (‘route-relevant’ information’). On the other hand, participants could search based on the immediate connections between nodes (Chrastil & Warren, 2014), causing them to depend on local information that is irrelevant to the route connecting the node to the target (‘route-irrelevant’ information). For instance, if participants started their search from nodes with a higher number of immediate connections, we would expect that their response times would increase as they searched irrelevant paths.

**Figure 4.**
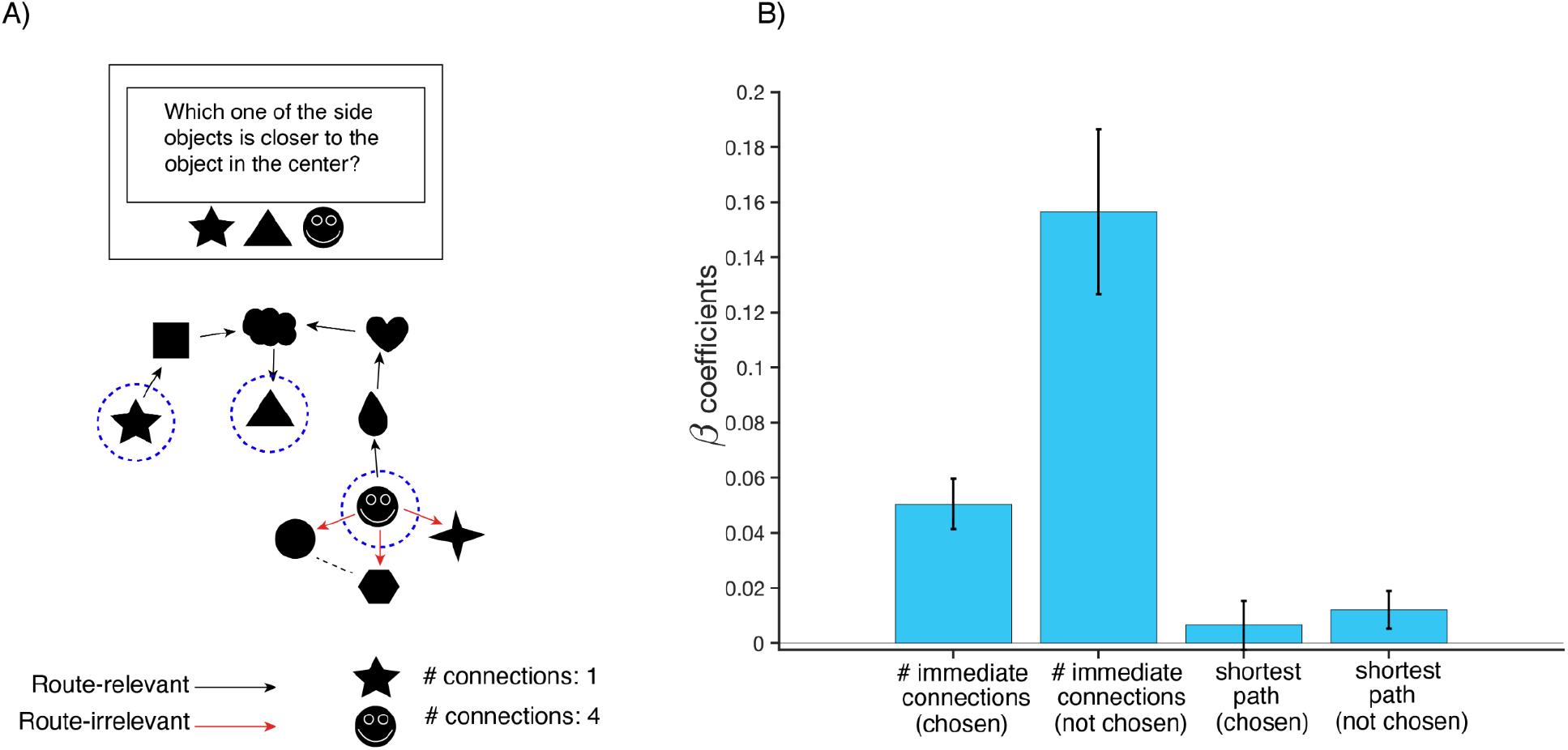
Participants incorporate irrelevant information into their graph search. **A)** If participants are using a global representation for search, they should be mostly sensitive to route-relevant information about the node-target distance. If participants are using local information for their search, they should be more sensitive to the number of route-irrelevant connections to the queried nodes. **B)** Regression coefficients show that the number of first degree connections is a stronger prediction of search time than the node-target distance, consistent with local search.

To test these two possibilities, we regressed participants’ judgment RTon the distance between the target and each of the queried nodes (as an index of route-relevant search), as well as on the number of immediate connections (degrees) of each of the queried nodes (as an index of route-irrelevant search). We found that judgment RTs were only weakly influenced by the distance between the target and both the chosen node (*β*=0.008, *p*=.33) and the unchosen node (*β* =0.01, *p*=.055) (Figure 4B). By contrast, RTs were significantly slower the more immediate connections either of the queried nodes had (chosen node: *β*=0.05, *p*=5.44e-08; unchosen node: *β*=0.15 *p*=7.78e-07). We further found that a model with both degree and distance information predicted reaction times better than one with only distance information (ΔAIC=122). These results support an interpretation whereby participants performed an extensive search of the graph, influenced by (route-irrelevant) local information about the queried nodes’ immediate connections.

After the judgment phase, we tested whether participants were also able to explicitly reconstruct this graph. On average, participants generated graphs that matched the true graph along two key metrics. First, these graphs successfully captured when two nodes were connected (i.e., formed an edge in the graph; *Figure 5A*, Mean Edge Accuracy = 84%, SD = 4%). Their overall accuracy at identifying these edges was substantially higher than would be expected by chance (e.g., if participants had configured the nodes at random; *Figure S1*).

**Figure 5.**
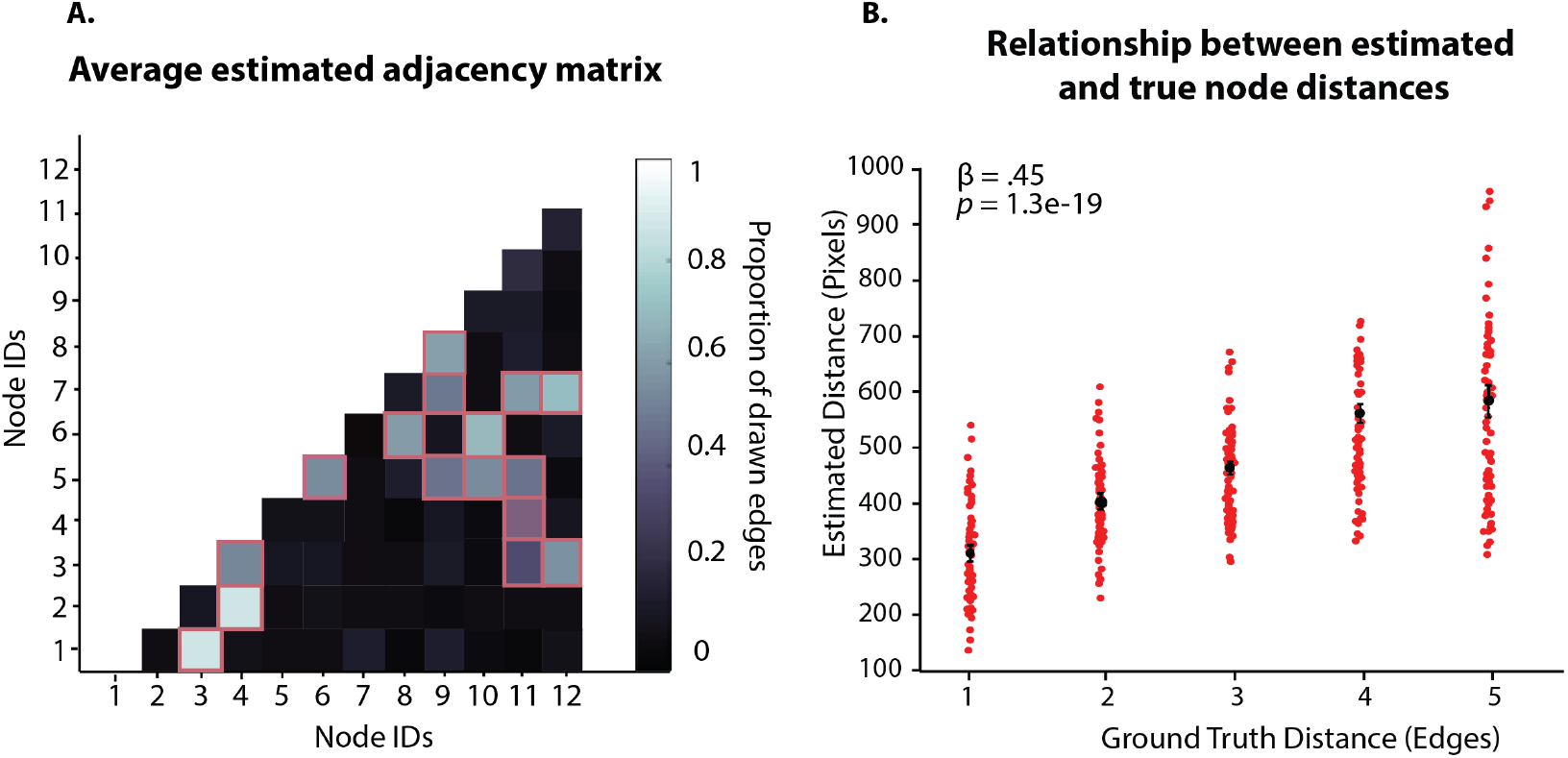
**A)** The proportion of drawn edges forming pairwise node connections in the graph. Lighter color indicates a higher proportion of connections. The squares outlined in orange correspond to the ground truth connections in the graph. Lighter color of fields outlined in orange indicates that the participants were more likely to draw edges between the nodes which are actually connected in the graph. **B)** Correlation between the ground truth distances (shortest paths in the graph), and pixel-based distances of recovered graphs.

Performance along this *adjacency* metric shows that participants were able to correctly identify the edges of the underlying graph, and therefore that they generally knew which nodes were connected to which other nodes. We also generated a second metric that examined the degree to which participants were able to also capture the relative *distances* between nodes in the graph (e.g., whether two connected nodes are close or far apart within the graph). In other words, when recreating the graph, to what degree was the Euclidean distance between objects placed on the screen (measured in pixels) representative of the true distance between the nodes in the underlying graph (measured in terms of the number of intervening edges/the shortest path). We found a significant correlation between these distance matrices (*Figure 5B*, *β* = .45, *t(76)* = 12.4, *p* = 1.3e-19, Mean(SD) Spearman *r* = .40(.25)). This relationship between estimated and true graph distance held even when controlling for the constructed adjacency matrix (*β* = .22, *t(76)* = 8.36, *p* = 1.2e-11, *Table S2*), suggesting that this distance metric captured structure inference ability over and above participants’ adjacency reports.

Our novel paradigm thus provides evidence that people are able to infer the structure of a graph based on experiences with disjoint edges from the graph, with neither explicit instruction of the graph’s existence nor a task goal that encourages them to learn this structure. We demonstrate this across six different measures: four from the relative distance judgment task (overall judgment accuracy, judgment accuracy by relative distance, search time by total distance, and search time by relative distance), and two measures from the graph reconstruction task (adjacency accuracy and the correlation between estimated (Euclidean) and actual (shortest path) graph distances). These six measures were highly correlated with one another across individuals (*Figure S2*), suggesting that they reflect a common underlying dimension of individual differences in structure inference ability. Indeed, a principal component analysis (PCA) demonstrated that a single component could capture 72% of the variance across these measures, and was the only substantive component that emerged in this analysis (all other eigenvalues < 0.85; *Figure 6B*). To examine the relationship between individual differences in structure inference ability and model-based planning, we therefore focused our analyses on variability in scores on this singular structure inference PC.

**Figure 6.**
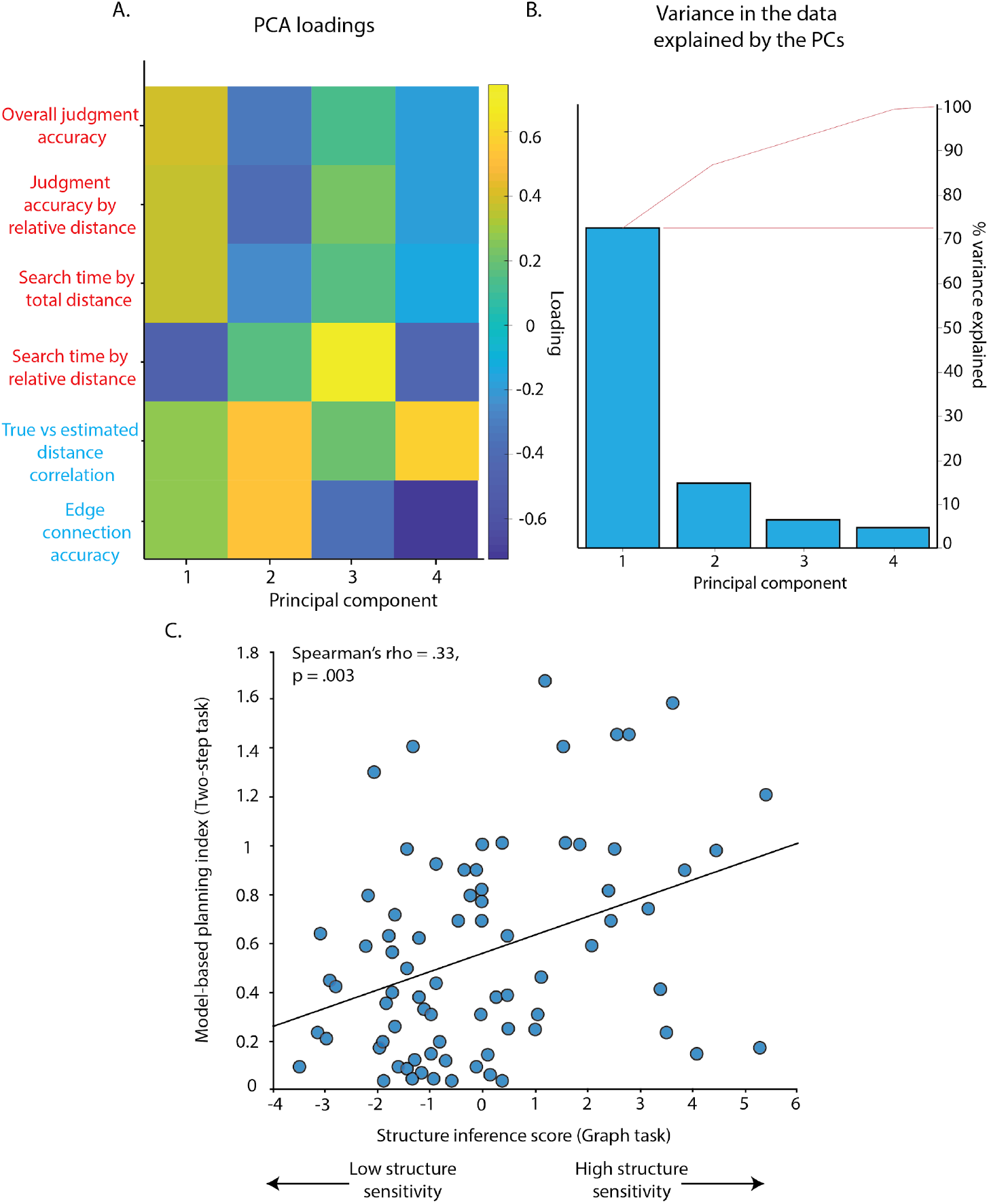
Plot A shows the PC loadings on all 6 graph-task measures (red = relative distance judgment task measures; blue = graph reconstruction measures). First component loads on all 6 measures, whereas the second component is selective for graph reconstruction measures. Eigenvalue of the first component is 4.33, and the second component is 0.85. Plot B shows the percentage of variance captured by different components. The first component (structure inference ability PC) captures the majority of variance (72%). Plot C shows that latent factor capturing structure inference ability (PCA score) is positively correlated with model-based planning (model-based weights β^*MB*^) across individuals.

### Better structure inference ability is associated with greater use of model-based planning

To measure individual differences in model-based planning, participants also performed a well-characterized assay of model-based planning, the two-step task (*Figure 1*, Daw et al., 2011; Gillan, Otto, Phelps, & Daw, 2015; Decker et al., 2016; Doll et al., 2015; Otto et al., 2013). In this task, participants make decisions at two stages that are connected by a probabilistic transition. To reach the best possible outcome on a given trial, participants must consider this underlying transition structure (e.g., engage in model-based planning). Previous studies show that choices in this task reflect a mixture of model-free and model-based forms of decision-making, indexing the degree to which participants choose actions based only on recent reward (*model-free*) or based additionally on a consideration of task structure (e.g., transition probabilities; *model-based*). We replicate this average pattern of behavior in our own data (model-free index: *β* = .64, *t*(76) = 14.38, *p* = 3e-05, model-based index: *β* =.42, *t(76)* = 9.17, *p* = 1.5e-20; *Figure 1C*, Table S1).

As in previous research, we also found that participants varied in their use of model-based planning on this task (e.g., as estimated by their model-based indices from the logistic regression). We predicted that these individual differences in model-based planning would be associated with individual differences in our index of structure inference ability, which we tested by examining whether there was a significant interaction between our structure inference PC and the model-based planning index in predicting first-stage choices (following previous work; Gillan et al, 2016). Consistent with our prediction, we found that participants who demonstrated better structure inference were also more likely to engage in model-based planning (*β* = .17, *t(76)* =3.97, *p* = .00008, *Figure 6C*, *Table S5*). This correlation held when using an alternate estimate of model-based planning, based on an RL model of the two-step task (*Figure 6C*, *R*^2^= .13, Spearman *r* = .33, *p* = .003; *Table S4*).

Follow-up analyses confirmed that structure inference ability was specifically associated with model-based planning and not other aspects of two-step task performance. We controlled for individual differences in model-free strategy use and stay bias (perseveration), neither of which were associated with our structural inference index (model-free strategy use: *β* = −.04, *t(76)* =−.17, *p* = .86; perseveration: *β* = −.11, *t(76)* =−.80, *p* = .42, *Table S3*). We also tested whether the relationship between structure inference and model-based planning was mediated by individual differences in model-based *learning*. That is, while the structure of the two-step task is relatively simple (each of two nodes transitioning to two other nodes), and participants are made aware of the potential links between these nodes in advance (though not of the actual transition likelihoods), it could be that people who are generally worse at inferring graph structure are less likely to engage in model-based planning because they failed to learn the transition structure of the two-step task. Previous work has measured such individual differences in two-step task transition learning using a separate behavioral index: response times for Stage 2 decisions (following the transition from Stage 1). Overall, participants have been shown to respond slower in Stage 2 if they just experienced a rare transition rather than a common one (Decker et al, 2016), a finding that we replicate in our own data (*β* = −.01, *t(76)* =3.54, *p* = .006). However, this effect depends on having learned which transitions are rare and common, and therefore individual differences in the strength of this effect have been used to index individual differences in transition learning. Unlike model-based *planning*, this implicit index of model-based *learning* (post-rare transition slowing) was *not* significantly associated with structure inference ability (*β* =.06, *t(76)* =.11, *p* = .57, *Table S3*). Moreover, controlling for model-based learning, the relationship between structure inference and model-based planning remained significant (*β* = .49, *t(76)* =2.19, *p* = .03, *Table S3*).

For completeness, we also tested whether the relationship between model-based planning and structure inference ability was specific to a subset of our structure inference metrics, but did not find that this was the case. Model-based planning was separately correlated with each of our structure inference metrics (all |*β*| > .12, all *p* < .007; *Table S5*).

### The relationship between MB-planning and structure inference ability is not accounted for by a single domain-general factor

While our results support a relationship between model-based planning and structure inference ability, they leave open the question of how direct this link is. For instance, the relationship between these variables could partially reflect their shared variance with individual differences in domain-general cognitive functions related to motivation, attention, and/or reasoning ability. Indeed, previous work has shown that MB reasoning correlates with some measures of general intelligence (Maran et al., 2020) and working memory capacity (Otto et al., 2013). To account for the influence of any such third variables, we compiled all available task performance measures and tested whether variability along these would be explained by a single dominant factor that linked our measures of interest (e.g., structure inference ability) with measures that reflect more general motivational and attentional processes (e.g., performance on our simple rotation discrimination task).

A PCA across 10 task variables (*Figure S4*) revealed that variability in performance on our attention check was in fact related to individual differences in other task variables that may have indexed motivation (e.g., degree of perseveration and reward sensitivity on the two-step task). However, this motivation-related factor was distinct from a factor that carried variance shared across all of our structure inference variables (e.g., judgment performance and edge estimation accuracy; *Figure S4 B,C*). This two-factor model suggests that our structure inference index measures a specific set of cognitive processes that are separable from ones that potentially relate to reward sensitivity and motivation. Importantly, our key ‘structure inference’ factor correlated with our MB measure even when partialling out this ‘motivation’ factor (partial *r(75)* = .29, *p* = 0.0096). This suggests that the relationship between MB and structure inference ability reflects processes over and above those that influenced general attentiveness across our tasks.

## Discussion

### Structure inference - general discussion

Acquiring structure representations is one of the most prominent examples of the robustness of human learning. People are able to learn the representations of their environments over the course of several trials, and leverage these representations in the service of problem solving. For instance, representations are thought to enable transfer of behavioral strategies between environments with similar structure. It is these benefits that distinguish humans from artificial agents, which currently fail to exhibit the flexibility commonly afforded by structure representations (Sutton et al., 1999).

Despite the evident benefits of structure representations, relatively little is known about how people infer/learn these structures in the first place. There is evidence that individuals are able to parse temporal streams of evidence into clusters, and use these temporal associations to infer relationships across states (Schapiro et al., 2012; Schapiro et al., 2016). The premise of such learning is that individuals are exposed to the full sequence of states in temporal order, and that such exposure enables individuals to extract statistical properties that support structure inference. However, it is also known that individuals are able to make judgments regarding states without ever experiencing transition between these states (Bunsey and Eichenbaum, 1996). This suggests that in addition to developing the structure representation based on statistical properties of temporal sequences, individuals are also capable of doing so by integrating across separate experiences. Therefore, it is essential to probe structure inference using a task that requires participants to carry out inferential processes based on disjoint information in order to piece together an overall structure. Our novel task represents an effort to capture this aspect of structure inference, and also provides additional evidence that individuals are biased to make inferences about structures, even when not told that there is one (Collins & Frank, 2016).

### Does structure inference constrain goal-directed behavior?

In addition to solving simple navigation problems, individuals can use underlying structures to perform goal-directed planning, which is critical for adaptive human behavior and long-term achievement across life domains. Successful goal-directed planning entails both (1) inferring the structure of one’s environment (structure inference) and (2) deploying that structure to maximize reward (model-based decision-making), yet relatively little is known about how one’s ability to do one of these relates to their ability to do the other. Combining six performance measures across our novel set of tasks, we characterized a dimension of structure inference ability, and showed that individual differences in this estimate of structure inference correlate with individual differences in a well-characterized index of model-based planning (based on performance on the two-step task). We show that this association between structure inference and two-step task performance is specific to model-based planning rather than generalizing to other performance metrics, including perseveration, model-free decision-making, and a proxy for one’s ability to learn transitions in that task. These results demonstrate that memory and decision-making share core cognitive substrates, with these connections across literatures potentially informing algorithmic models of knowledge-driven planning and decision-making. Given this initial validation, our tasks hold promise for further bridging these lines of research to examine goal-directed navigation towards a reward (discussed below).

Our work bridges previous research on structure learning and model-based planning, and addresses an important gap in these earlier studies. In particular, previous research on model-based planning has only been able to examine how people learn about and navigate internal models with limited nodes/connections (e.g., a single transition in the two-step task; Daw et al. 2011) and/or with transitions that are experienced sequentially in time (Bornstein & Daw, 2013; Doll et al., 2015). Using our novel task, we were able to examine how this inference process occurred for a relatively more complex graph structure that participants learned based on disjoint experiences with individual node pairs. In doing so, we were able to establish that individuals are able to perform structure inference under such conditions and to exploit individual differences in their inferential abilities on these tasks to link the underlying cognitive processes to variability in model-based planning within simpler environments.

## Limitations

### Structure inference task

An important limitation of our study is that we do not have measures of structure inference taken prior to the judgment phase of the task (Phase 2). As a result, our measures of structure inference can reflect both (a) a participant’s ability to infer the graph structure during the latent exposure phase (Phase 1) and (b) their ability to infer this graph structure in subsequent phases based on their initial exposure (possibly via directed search through these learned associations). Therefore, to tease apart the learning processes during the initial exposure vs. additional build-up of the structure during reconstruction, one would need to implement assessment of online learning during the initial exposure, potentially by implementing occasional probes about the structure.

Furthermore, while our study was able to provide an in-depth examination of how participants learn about the structure of the environment, it did so in the context of a fairly small structure (twelve nodes total) with deterministic transitions. This limits our ability to estimate how well our participants would be able to learn and traverse wider and more complex structures. One feature of many real-world structures that facilitates learning is temporal contiguity between nodes, something that we explicitly sought to control in order to isolate structure inference from sequence learning.

### Relating structure inference and model-based planning

We have demonstrated an initial validation of the potential correspondence between structure inference and model-based planning by revealing a correlation between our compound measure of structure inference and indices of model-based planning. However, our inability to clearly dissociate between online learning and memory-based inference (driven by exposure during later phases) prevents us from testing which one of these, if not both, relates to model-based planning.

We are also unable to completely rule out the possibility that this correlation was partly driven by domain-general factors like motivation, executive function, working memory, or fluid intelligence, factors that have been shown to correlate with model-based performance (Maran et al., 2020; Otto et al., 2013). While our PCA analyses provide evidence that plausible indices of task engagement load on a separate component from putative measures of structure inference and MB, it is important to confirm this with more explicit measures of each of these domain-general constructs.

## Future directions

### Neural mechanisms of structure inference

The observation that humans are able to rapidly learn complex graph structures without prior awareness raises several questions for further investigation. A particularly valuable direction for future research will be to investigate the neural mechanisms by which individuals infer the structure of our graph: Are these links inferred at encoding, during the learning task, or on-demand, at each trial during the judgment task? Evidence for both mechanisms was observed in a previous study that investigated transitive inferences across pairs of words in humans undergoing intracranial EEG (Reber et al. 2016). The authors reported that successful later inferences were predicted by hippocampal activity evident in both response-locked ERPs at test and stimulus-locked ERPs following encoding, providing support for multiple, hippocampally-centered mechanisms in the construction of inferences. This observation is consistent with a previously proposed distinction between prospective and retrospective integration in support of memory-guided decisions (Shohamy & Daw 2015; Ballard et al. 2019), and with recent work separately identifying a role for both encoding-time (Schapiro et al. 2013; Schapiro et al. 2016) and retrieval-time (Kӧster et al. 2018) computations in supporting similar inferential judgments. In both cases the judgments tested have been limited to a single step, though a more extensive capability was implied. Further work will be necessary to identify the relative contribution of each of these mechanisms to our task, in particular whether encoding and retrieval mechanisms are similarly useful in identifying extensive latent structure.

### Utilizing latent structures for transfer learning

A critical factor in the ability to infer latent structure might be the effective use of hypotheses about the structure of the task. Specifically, humans and other animals are capable of *transfer learning*, or applying a schema learned in one instance of a task to another. Because our novel graph task relies on structures that can be clustered into families of resemblance and permuted to varying degrees, it is well-suited to measure the extent of the ability to transfer across multiple instances. An important open question is to what degree these inferences are supported by general conceptual representations e.g. in PFC (Kumaran et al. 2012), structured basis representations in MTL cortex (Schapiro et al. 2016; Constantinescu et al. 2016; Behrens et al. 2018), or pre-generated associations cached in hippocampus proper (Collin et al. 2015). Because our task permits finely manipulating the relative information present in each kind of representation, it may be suitable for distinguishing involvement of each of these neural substrates.

### Further investigations of structure inference-MB planning relationship

Having provided initial evidence of the potential relationship between structure inference and model-based planning, future investigations could extend the task to incorporate planning for rewards directly, for instance by having participants learn about state-reward associations after (or in parallel with) learning the structure of state-state associations (Wimmer et al. 2012; Bornstein & Daw 2013). Examining how participants navigate this structure can provide important insight into how the structure of an internal model, and the relative distance between nodes in that model, modulates the utility of future rewards (Kurth-Nelson et al., 2015; Wimmer & Shohamy, 2012; Bornstein & Daw 2013) and the mental effort required to obtain those rewards (Kool & Botvinick., 2018, Shenhav et al., 2017). A similar design can also provide critical links to an expansive body of work on goal-directed navigation through space (Tolman, 1948; Tolman and Honzik, 1930; Tolman et al., 1946), including factors that influence individual differences in one’s success in such navigation tasks (Maguire et al., 2000). In addition to highlighting common and divergent mechanisms across these domains, such research can also point toward potential training regimens that can be used to improve goal-directed planning, building on recent work in this area (Lieder et al., 2019). Collectively, these tasks can be used to compare and contrast neural mechanisms that underpin learning, navigation, and planning over latent structures across spatial and non-spatial domains.

### Mechanistic underpinnings of model-based planning impairments

Our work also provides important methodological and mechanistic insight for research on goal-directed decision-making across the lifespan, and between healthy and clinical populations. For instance, prior work has demonstrated that model-based planning gets worse with advanced aging (Eppinger et al., 2013) and negatively scales with behaviors indicating compulsive symptoms (Gillan et al., 2016). However, it remains unclear to what extent such impairments arise from deficits in inference (e.g., an inability to acquire and/or retain an internal model of the associative structure of one’s environment) and/or deficits in the ability/motivation to search that model to determine the best course of action (e.g., due to working memory demands and attendant mental effort costs). A better understanding of the mechanisms at the intersection of these processes will provide critical insight into the nature of goal-directed planning, and the factors that determine one’s ability to achieve those goals.

## Acknowledgments

This work was supported by a Center of Biomedical Research Excellence grant (P20GM103645) from the National Institute of General Medical Sciences (A.S.). The authors are grateful to Anna Schapiro, Avinash Vaidya, Matthew Nassar and Peter Dayan for helpful discussions.

## Supplemental Figures

**Figure S1.**
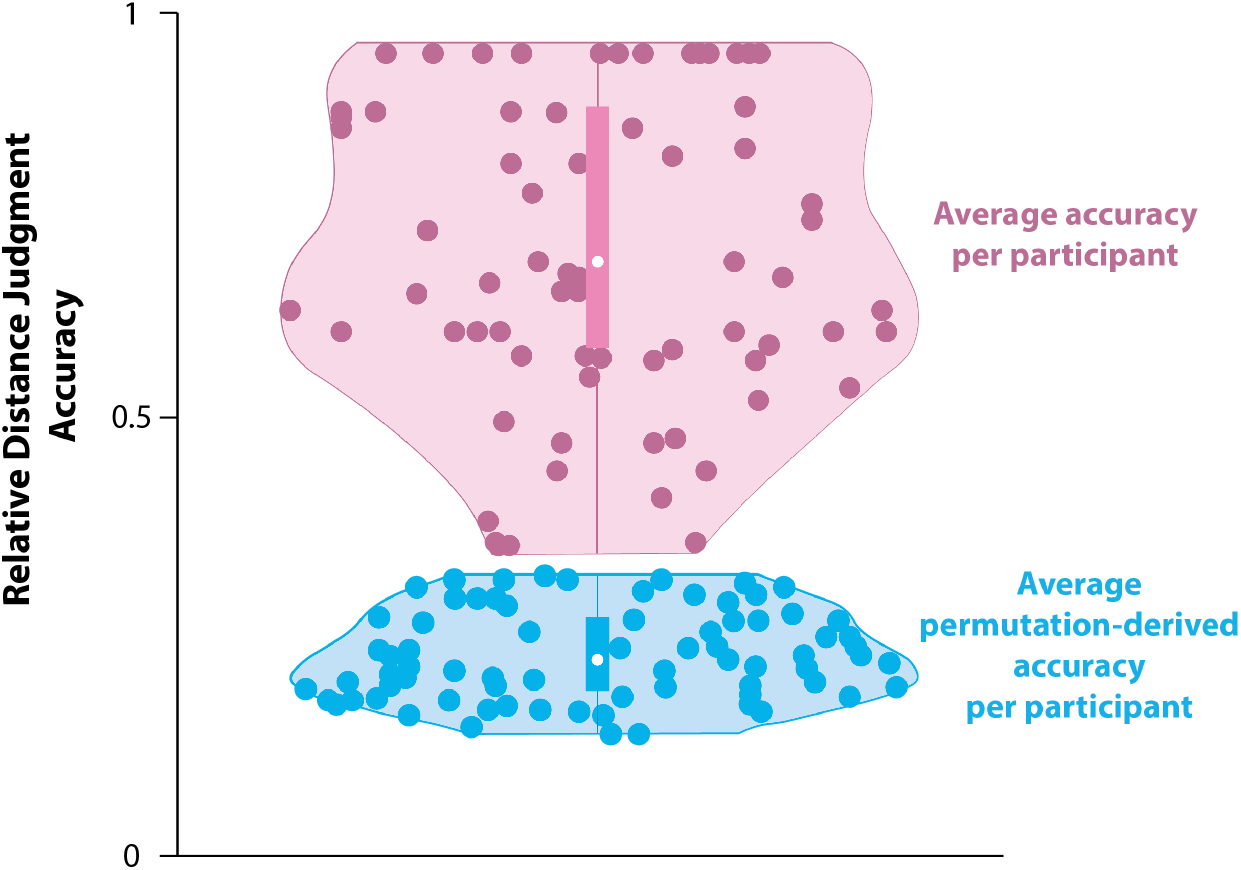
Distribution of (1) participant accuracy on graph reconstruction task (the percentage of correctly identified edges) and (2) chance level (permutation-derived) accuracy for each subject. We estimated chance accuracy for each subject by randomly shuffling the graph 1000 times (such that connections between nodes differ from those in the ground truth graph), evaluating the accuracy of connected edges based on the adjacency matrix of the shuffled graph and averaging accuracy values in each subject. The edge connections based on the shuffled graph are a proxy of how accurate subjects would be if they were guessing at random while drawing the edges between nodes. Our results show that participants’ edge connection accuracy evaluated based on the ground truth adjacency matrix is significantly greater than the chance accuracy based on the randomly shuffled graphs, confirming that on average participants were not randomly configuring the edges.

**Figure S2.**
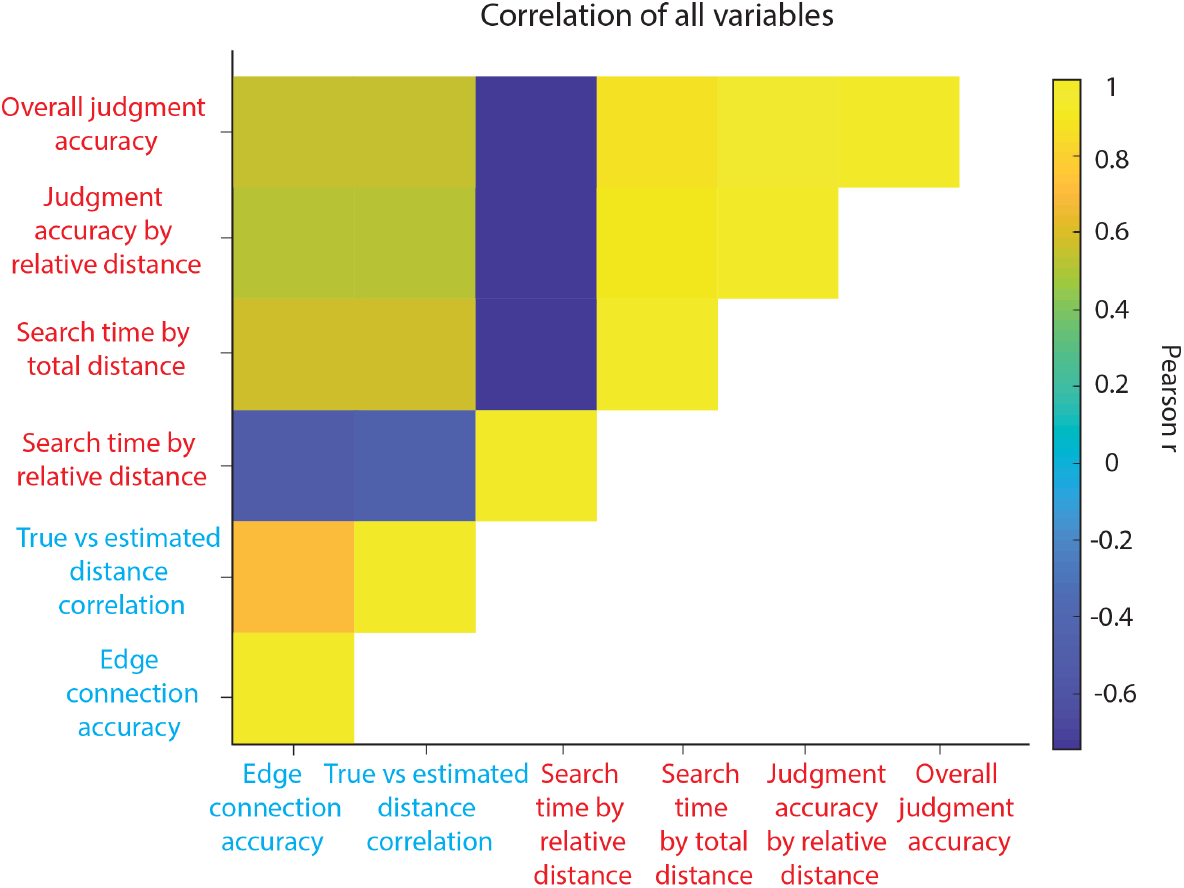
Pairwise correlations between all graph-task measures.

**Figure S3.**
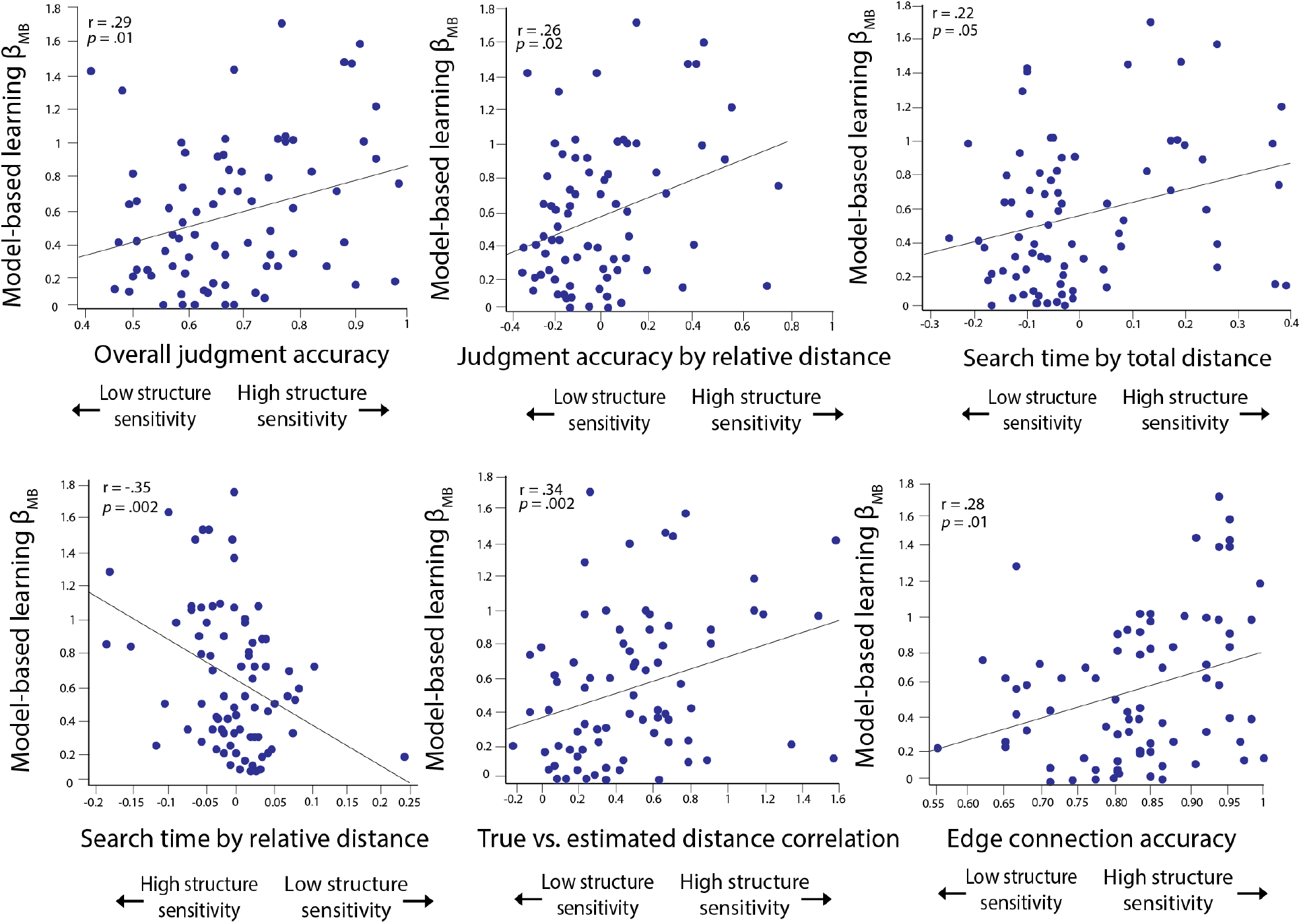
Model-based learning is associated with individual structure inference measures. The y-axis corresponds to the fit value of the model-based weighting parameter *β*_MB_. The x-axis in the above figures plots the following: overall judgment accuracy, judgment accuracy by relative distance, search time by total distance, search time by relative distance, true vs. estimated distance correlation, edge connection accuracy.

**Figure S4.**
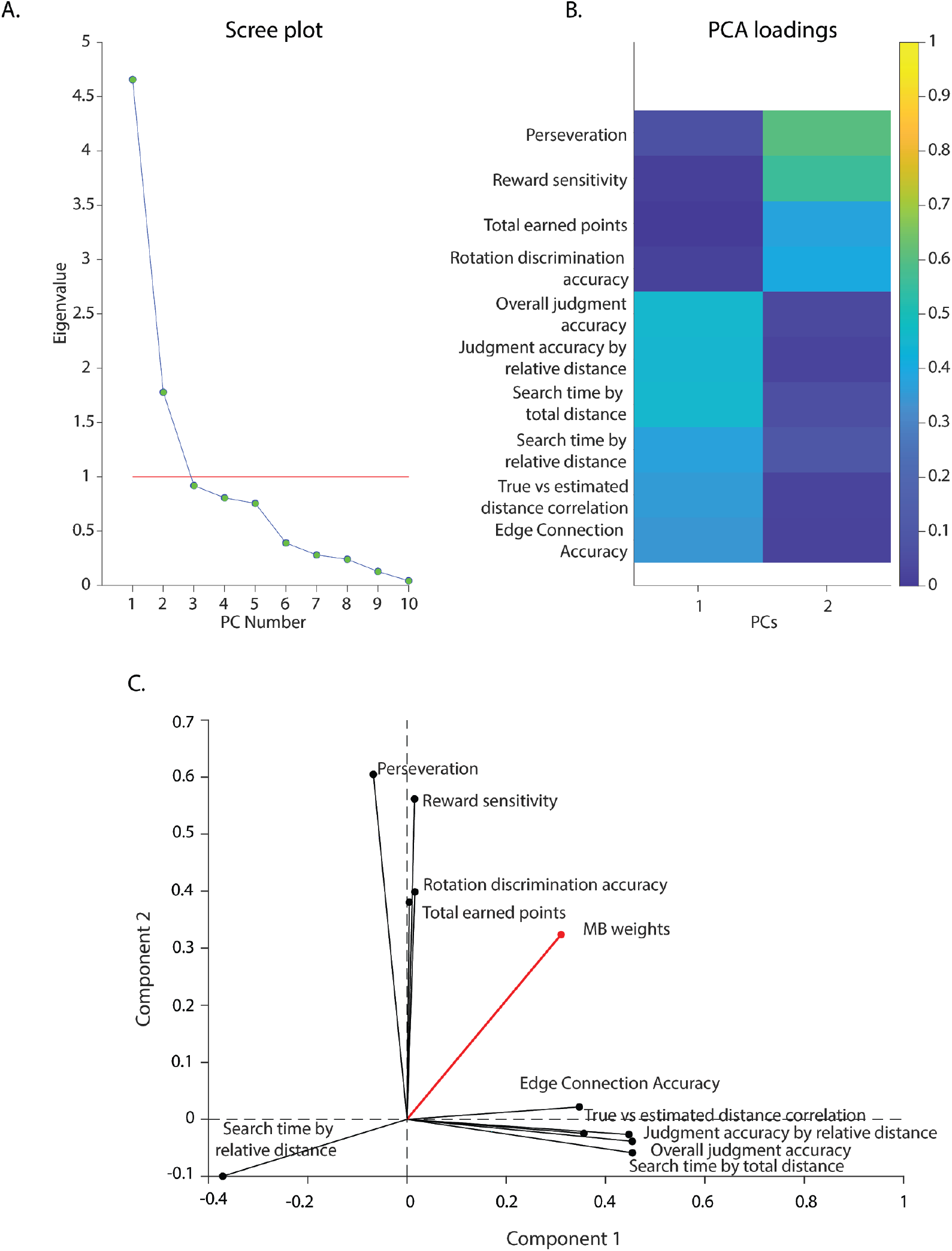
Exploratory PCA on individual differences. Our exploratory PCA (promax rotated) revealed that the data supported a two component model (A), with factors putatively related to structure learning (Component 1) and motivation (Component 2; B). (C) Our model-based index loaded on to both of these components.

## Supplemental Tables

**Table S1.** Modeling 1st-stage choices in the RL task as a function of model-free and model-based learning. Model statistics refer to the coefficients of the fixed main effect of reward, transition type and the reward-by-transition type interaction from the following model: *Stay ~ Reward × Transition type + (1 + Reward × Transition type | Participant)*. Here (SE) indicates the standard error of the mean (*p<.05; **p<.01; ***p<.001). Participants exhibit a mixture of model-free and model-based strategies, as shown by the significant main effect of reward and a reward-by-transition type interaction.

| | $\beta$(SE) | $T$ | DF | $p$ |
| --- | --- | --- | --- | --- |
| Transition type | .10(.02) | 3.9 | 73 | .0001*** |
| Reward | .64(.04) | 14.38 | 73 | 2.9e-05*** |
| Transition type *<br>Reward | .42(.04) | 9.17 | 73 | 1.5e-20*** |

**Table S2.** Predicting ground truth distance with Euclidean distance, controlling for reported shortest path. The Euclidean distance (estimated distance) predicts the ground truth distance over and above the reported shortest path. This suggests that the estimated distance model was not predictive of the ground truth simply as a function of edge connections participants drew during the reconstruction. DF here refers to Satterthwaite degrees of freedom approximation. (*p<.05; **p<.01; ***p<.001).

| | $\beta$(SE) | $T$ | DF | $p$ |
| --- | --- | --- | --- | --- |
| Estimated<br>shortest path | .42(.05) | 7.81 | 74 | 5.63e-11*** |
| Euclidean/<br>distance | .22(.02) | 8.36 | 74 | 1.22e-11*** |

**Table S3.**
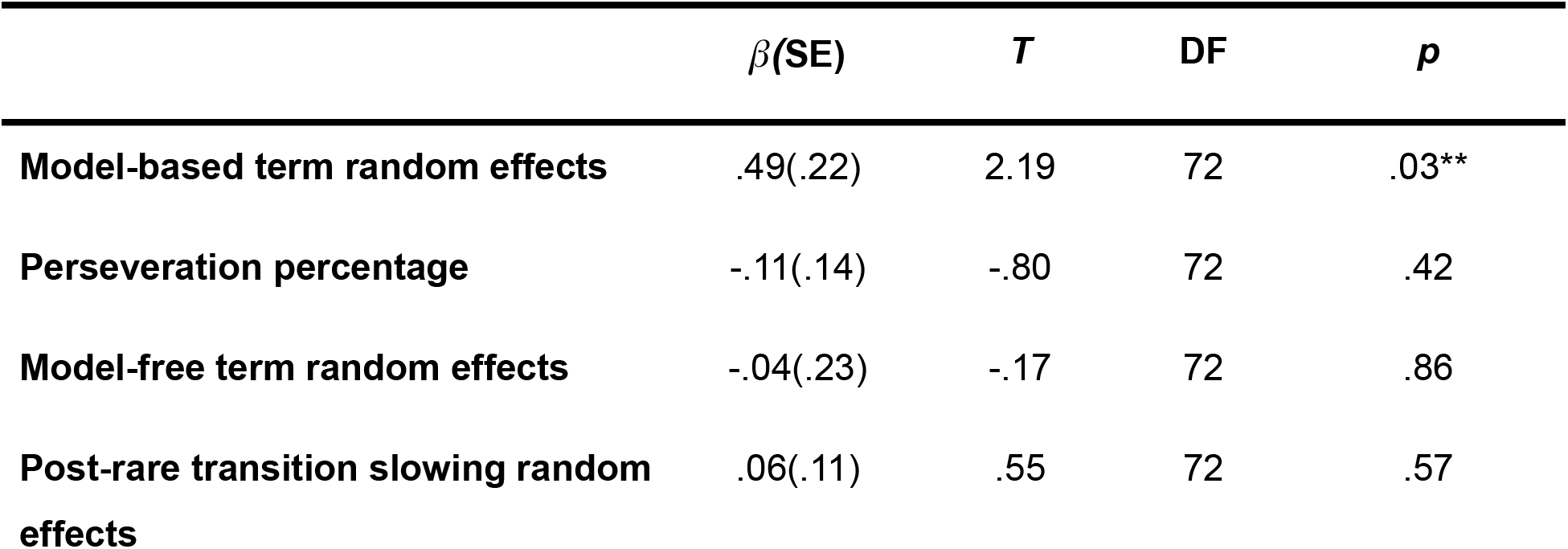
Robust linear regression model predicting PC scores (latent component indexing structure inference) using different indices from the two-step task (percentage of perseverative response, model-free random effects, model-based random effects and post-rare transition slowing). Index of model-based planning is selectively significantly associated with the measure of structure inference. The beta coefficients here are estimated effect coefficients, SE is standard error of the mean. DF refers to error degrees of freedom. Positive terms indicate positive association with the structure inference measure (*p<.05; **p<.01; ***p<.001).

| | $\beta$(SE) | *T* | DF | *p* |
| --- | --- | --- | --- | --- |
| **Model-based term random effects** | .49(.22) | 2.19 | 72 | .03** |
| **Perseveration percentage** | -.11(.14) | -.80 | 72 | .42 |
| **Model-free term random effects** | -.04(.23) | -.17 | 72 | .86 |
| **Post-rare transition slowing random effects** | .06(.11) | .55 | 72 | .57 |

**Table S4.**
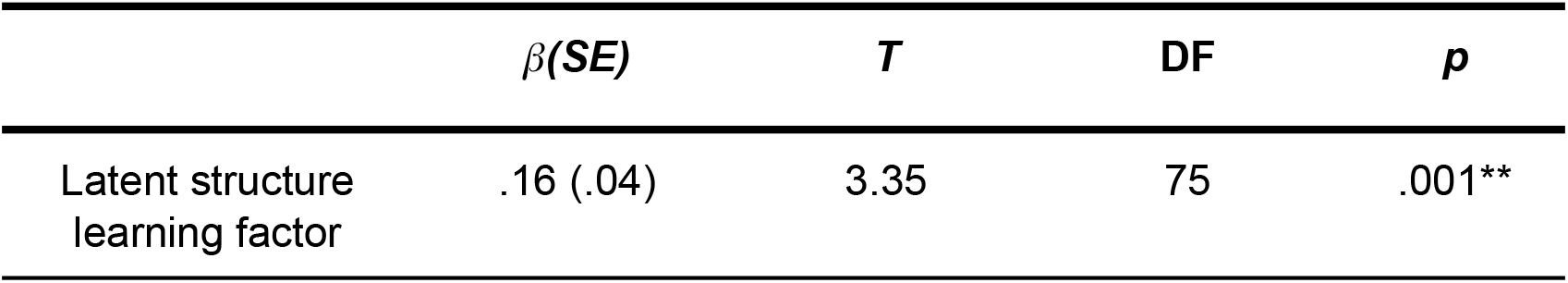
Model-based weights from the computational model. The results from the robust linear regression model (*RL Model-based weights ~ 1 + structure inference score*). The predictor in the model was z-scored. The results show that high PC scores (indexing high structure inference performance) predict increased model-based planning. (*p<.05; **p<.01; ***p<.001).

**Table S5.** Modeling 1st-stage choices in the RL task as a function of structure inference ability and model-based planning. Each row reflects the results from an independent analysis where each covariate (z-transformed) was entered as Z in the following model: *Stay ~ Reward × Transition × Z + (1 + Reward × Transition | Participant)*. Model statistics refer to the coefficient of the fixed-effects interaction: *Reward × Transition × Z*. Positive values indicate an association with increased model-based planning Covariates with positive values are associated with increased model-based learning (except for the search time by relative distance effect). Here (SE) indicates the standard error of the mean. (*p<.05; **p<.01; ***p<.001).

| Structure inference measure | $\beta$ (SE) | *T* | DF | *p* |
| --- | --- | --- | --- | --- |
| Overall distance judgment accuracy | .14 (.05) | 3.16 | 72 | .001** |
| Judgment accuracy by relative distance | .16 (.04) | 3.60 | 72 | .0003*** |
| Search time by total distance | .13(.04) | 2.95 | 72 | .003** |
| Search time by relative distance | -.16 (.04) | -3.69 | 72 | .0002*** |
| True vs estimated distance correlation | .18 (.04) | 3.98 | 72 | .00008*** |
| Edge connection accuracy | .12 (.04) | 2.67 | 72 | .007** |
| Latent structure learning factor | .17(.04) | 3.97 | 72 | .00008*** |

**Table S6.** Graph task performance variables index.

| Performance measure names | Definition |
| --- | --- |
| Overall judgment accuracy | Proportion of trials on which participants correctly identified the closer node |
| Judgment accuracy by relative distance | The effect of distance discrepancy between the option nodes and the target node on accuracy. Greater distance discrepancy should lead to higher accuracy. |
| Search time by total distance | The effect of overall distance of two option nodes to the target on response times. The greater overall distance should lead to longer response times, since participants would need to perform a longer 'search' to make their response. |
| Search time by relative distance | The effect of distance discrepancy between the option nodes and the target node on |
|  | response times. Greater distance discrepancy should lead to faster responses. |
| True vs estimated distance correlation | In the edge reconstruction task, we correlated the pairwise node distance given by 1) number of pixels between two node images in the reconstructed graph, and 2) the pairwise shortest path (number of edges) between all nodes in the underlying graph |
| Edge connection accuracy | Proportion of correctly identified edges, based on the comparison of node connections in the recovered graph to ground truth adjacency matrix |

